# Reticulocalbin 3 is Involved in Postnatal Tendon Development by Regulating Collagen Fibrillogenesis and Cellular Maturation

**DOI:** 10.1101/2020.09.25.313098

**Authors:** Na Rae Park, Snehal Shetye, Douglas R. Keene, Sara Tufa, David M. Hudson, Marilyn Archer, Louis J Soslowsky, Nathaniel A. Dyment, Kyu Sang Joeng

## Abstract

Tendon plays a critical role in the joint movement by transmitting force from muscle to bone. This transmission of force is facilitated by its specialized structure, which consists of highly aligned extracellular matrix consisting predominantly of type I collagen. Tenocytes, fibroblast-like tendon cells residing between the parallel collagen fibers, regulate this specialized tendon matrix. Despite the importance of collagen structure and tenocyte function, the biological mechanisms regulating fibrillogenesis and tenocyte maturation are not well understood. Here we examine the function of Reticulocalbin 3 (Rcn3) in collagen fibrillogenesis and tenocyte maturation during postnatal tendon development using a genetic mouse model. Loss of Rcn3 in tendon caused decreased tendon thickness, abnormal tendon cell maturation, and decreased mechanical properties. Interestingly, Rcn3 deficient mice exhibited a smaller collagen fibril distribution and over-hydroxylation in C-telopeptide cross-linking lysine from α1(1) chain. Additionally, the proline 3-hydroxylation sites in type I collagen were also over-hydroxylated in Rcn3 deficient mice. Our data collectively suggest that Rcn3 is a pivotal regulator of collagen fibrillogenesis and tenocyte maturation during postnatal tendon development.

## Introduction

Tendons are critical for native function of the musculoskeletal system. They are primarily composed of type I collagen and individual collagen fibrils arrange themselves into the highly organized bundles that can resist high tensile forces^1^. Tendons undergo significant changes in tissue and cell morphology throughout postnatal development. Tendon structure becomes denser and more organized via matrix production and collagen fibrillogenesis. Cell density becomes lower as matrix production increases and cell proliferation decreases^2^. The shape of tendon cell nuclei transform from round to a short spindles, and eventually to a long, flat spindles^3^. Previous studies have shown that several signaling pathways and transcription factors are critical regulators of embryonic and postnatal tendon development, but the precise regulatory mechanisms for cellular maturation and collagen fibrillogenesis during postnatal tendon development are still not clear^4–11^.

Developmental stage of tendons is marked by an increase in the number of collagen fibrils deposited in the matrix^12^. However, postnatal stages are shown to increase the diameter of existing collagen fibrils rather than an increase in collagen fibril number^12^. This transition occurs in the rough endoplasmic reticulum and involves a complex process wherein small collagen fibrils undergo lateral fusion to form thicker fibrils^13^. Mutations in several genes that regulate this process, including Decorin, Fibromodulin, and Lumican can cause defects in tendon maturation^14–19^.

Collagens consist of repeating Gly-X-Y triplets. Procollagen biosynthesis occurs in the ER and has characteristic post-translational modifications, including hydroxylation and glycosylation. Two modifications of proline residues are known, prolyl 4-hydroxylation and prolyl 3-hydroxylation^13^. Almost all proline residues lying in the Y position are prolyl 4-hydroxylation. Thus prolyl 4-hydroxylation stabilizes the collagen triple helix because the melting temperature of the collagen triple helix is directly proportional to the prolyl 4-hydroxylation content^20,21^. Several tissue-specific prolyl 3-hydroxylation modification sites have been identified in tendon type I collagen, including α1(I) Pro986, α1(I) Pro707, α2(I) Pro707, and a C-terminal (GPP)n motif that are mainly modified by P3H2 and α1(I) Pro986 that is modified by P3H1^22^. The total prolyl 3-hydroxylation content in the adult Achilles tendon collagen was higher than that in fetal Achilles tendon collagen^23^. Several studies suggested that prolyl 3-hydroxylation is involved in collagen fibril assembly^22,24,25^.

Reticulocalbin 3 (Rcn3), a 45 kDa ER lumen protein, is a member of the CREC (Cab45/reticulocalbin/ERC45/calumenin) family of multiple EF-hand Ca^2+^-binding proteins, including Rcn1, ERC-55 (Rcn2), Rcn3, Cab45, and calumenin^26^. Proteomic studies have suggested that Rcn3 functions as a chaperone protein in the secretory pathway and has emerged as a new potential regulator of collagen production^27,28^. Recent mouse genetic studies have demonstrated that Rcn3 plays a critical role in perinatal lung maturation by regulating synthesis and secretion of surfactant and is involved in pulmonary injury remodeling via regulation of alveolar epithelial cell (AEC) apoptosis and ER stress^29,30^. Despite this emerging role of Rcn3 in the secretory pathway, the physiological function of Rcn3 in various tissues producing a large amount of extracellular matrix is not yet fully established.

In this study, we investigated the physiological function of Rcn3 in tendon by conditionally deleting Rcn3 in mouse tendon cells. Loss of Rcn3 caused impaired postnatal tendon maturation with markedly decreased tendon thickness. Consistently, structural properties were reduced in Rcn3 deficient mice but material properties were mostly unchanged. We further found that Rcn3 deficiency resulted in impaired collagen fibrillogenesis with over-modification of type 1 collagen and reduced fibril diameter distribution. Collectively, these findings, for the first time, establish the biological function of Rcn3 in postnatal tendon development and provide an exciting mouse genetic model for studying collagen fibrillogenesis and modification in tendon.

## Results

### Loss of Rcn3 caused decreased tendon thickness

To understand the function of Rcn3 in tendons, we generated *Scx-Cre; Rcn3*^*fl/fl*^ mice in which Rcn3 was specifically deleted in Scx-expressing cells that are abundant in tendons^31,32^. The growth of *Scx-Cre; Rcn3*^*fl/fl*^ mice was normal compared with wild-type mice, which was evident with normal body weight (Figure 1A). To access the knockout efficiency of Rcn3 in tendons, we performed immunohistochemistry analysis against Rcn3 (Figure 1B). Rcn3 was highly expressed in patellar and Achilles tendons in wild-type mice at the early postnatal stage, including P5, P21, and P30, but its expression was decreased at a later stage, P60 (Figure 1B). Immunohistochemistry analysis demonstrated that the deletion of Rcn3 in *Scx-Cre; Rcn3*^*fl/fl*^ tendons were efficient in both patellar and Achilles tendons throughout the postnatal maturation (Figure 1B). In the patellar tendon, we observed residual Rcn3 expression at P5 and P21, but the expression was not detected at P30 and P60. In the Achilles tendon, we found residual expression till P30, but the expression was not detected at P60. These data suggest that the deletion of Rcn3 is more efficient in the patellar tendon than the Achilles tendon.

**Figure 1.**
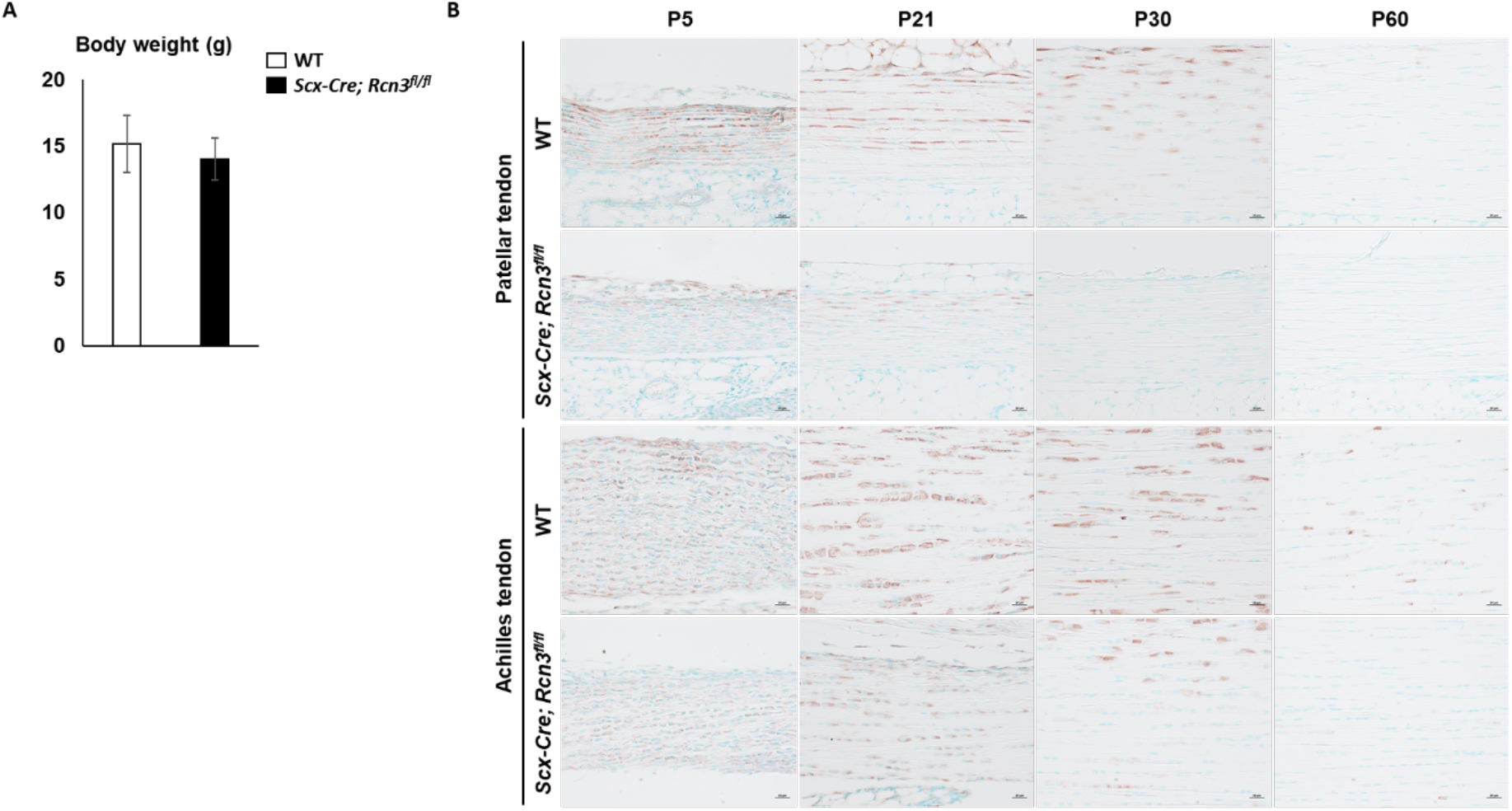
The weight of wild-type mice and *Scx-Cre; Rcn3*^*fl/fl*^ mice at P30 (A). Immunohistochemical analysis of Rcn3 on patellar and Achilles tendons from wild-type mice and *Scx-Cre; Rcn3*^*fl/fl*^ littermates during postnatal tendon maturation (B). (Brown color indicates Rcn3, Scale bar indicates 20µm (B) and n=5 (A))

We performed histological analyses on patellar and Achilles tendons to assess the tendon phenotypes in *Scx-Cre; Rcn3*^*fl/fl*^ mice. Although *Scx-Cre; Rcn3*^*fl/fl*^ mice had no difference in body weight compared to wild-type mice (Figure 1A), *Scx-Cre; Rcn3*^*fl/f*^ mice clearly displayed decreased tendon thickness at P30 compared to wild-type littermate in both patellar (Figure 2A) and Achilles tendon (Figure 2C). Our quantification results confirmed the 40% reduced thickness in the patellar tendon and 20% reduced thickness in the Achilles tendon (Figure 2B and 2D). We also found that the cellularity of tendons in *Scx-Cre; Rcn3*^*fl/fl*^ mice was higher than wild-type mice, with 3-fold and 1.5-fold increases in the patellar and Achilles tendons, respectively (Figure 2B and 2D). The more severe phenotypes in the patellar tendon are consistent with our deletion efficiency result (Figure 1B). These results collectively suggest that Rcn3 is critical for postnatal tendon development for both patellar and Achilles tendons.

**Figure 2.**
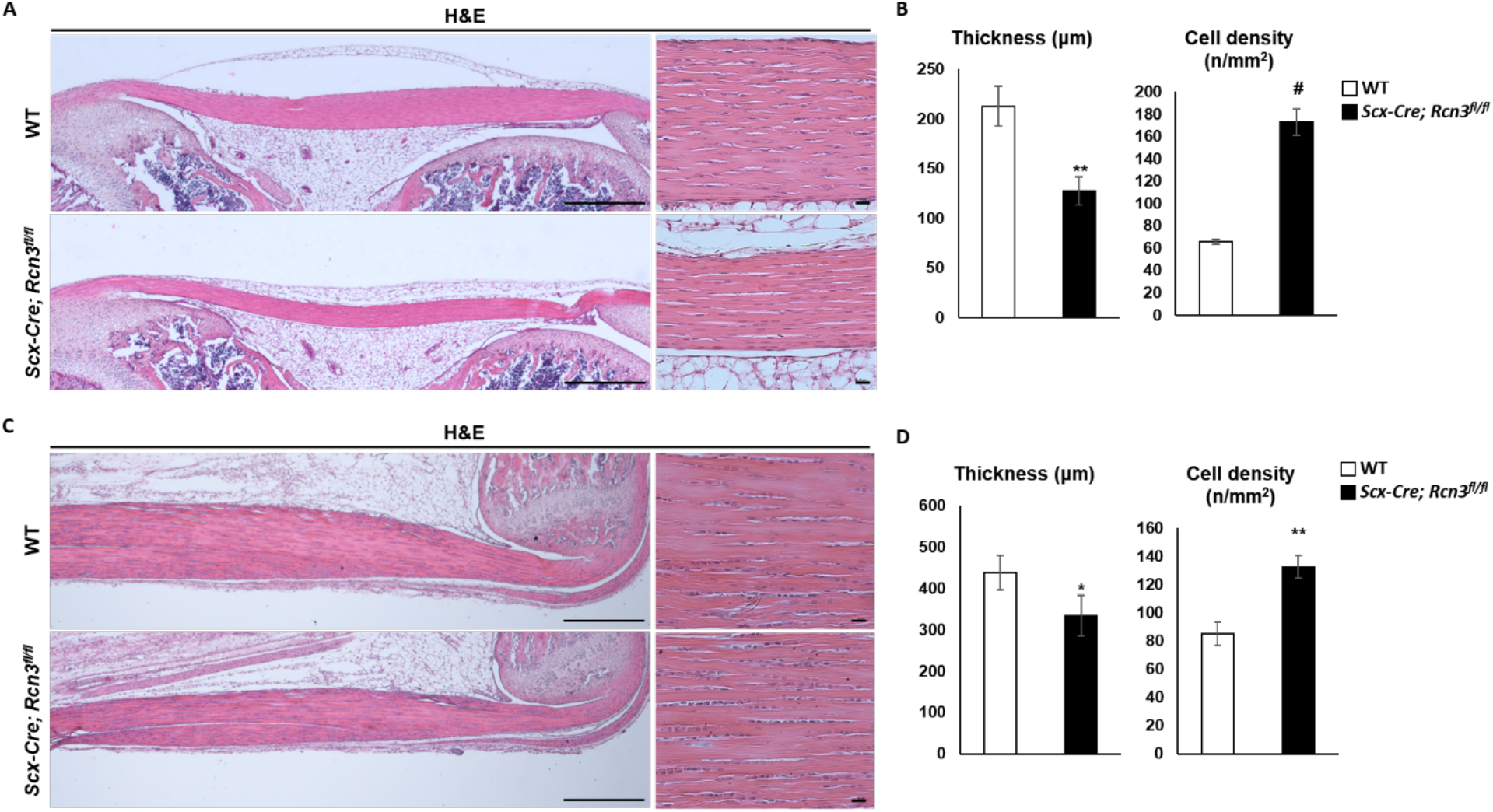
Histology of patellar and Achilles tendons from wild-type mice and *Scx-Cre; Rcn3*^*fl/fl*^ littermates during postnatal tendon development (A and C). The thickness and cell density of patellar tendon and Achilles tendon at P30 (B and D). (Scale bar indicates 500µm (A) and 20µm (B), * indicates *P*<0.05, ** indicates *P*<0.01 and ^#^ indicates *P*<0.001 between genotypes, n=3)

### Loss of Rcn3 reduced collagen fibril diameter

The decreased tendon thickness prompted us to examine collagen fibril ultrastructure because collagen fibrillogenesis is critical for tendon growth. We performed transmission electron microscopy (TEM) using patellar and Achilles tendons at two months of age to measure collagen fibril diameter. *Scx-Cre; Rcn3*^*fl/fl*^ mice exhibited decreased fibril diameter in both patellar (Figure 3A) and Achilles tendons (Figure 3C) compared with wild-type mice. Specifically, quantification results showed that the distribution of collagen fibril diameter in patellar tendon ranged from 20 to 210nm in wild-type mice, whereas *Scx-Cre; Rcn3*^*fl/fl*^ mice showed a narrower collagen fibril distribution (10 to 170nm) in the patellar tendon lacking the larger fibrils found in WT mice (Figure 3B). Consistent with the patellar tendon, the Achilles tendon also showed the same trend with a shifted distribution in the mutants lacking the larger fibrils (i.e., 190-260 nm) found in WT mice (Figure 3D). These results suggest that Rcn3 is required for proper collagen fibrillogenesis, and decreased tendon thickness could be due to the smaller diameter of collagen fibril in *Scx-Cre; Rcn3*^*fl/fl*^ mice.

**Figure 3.**
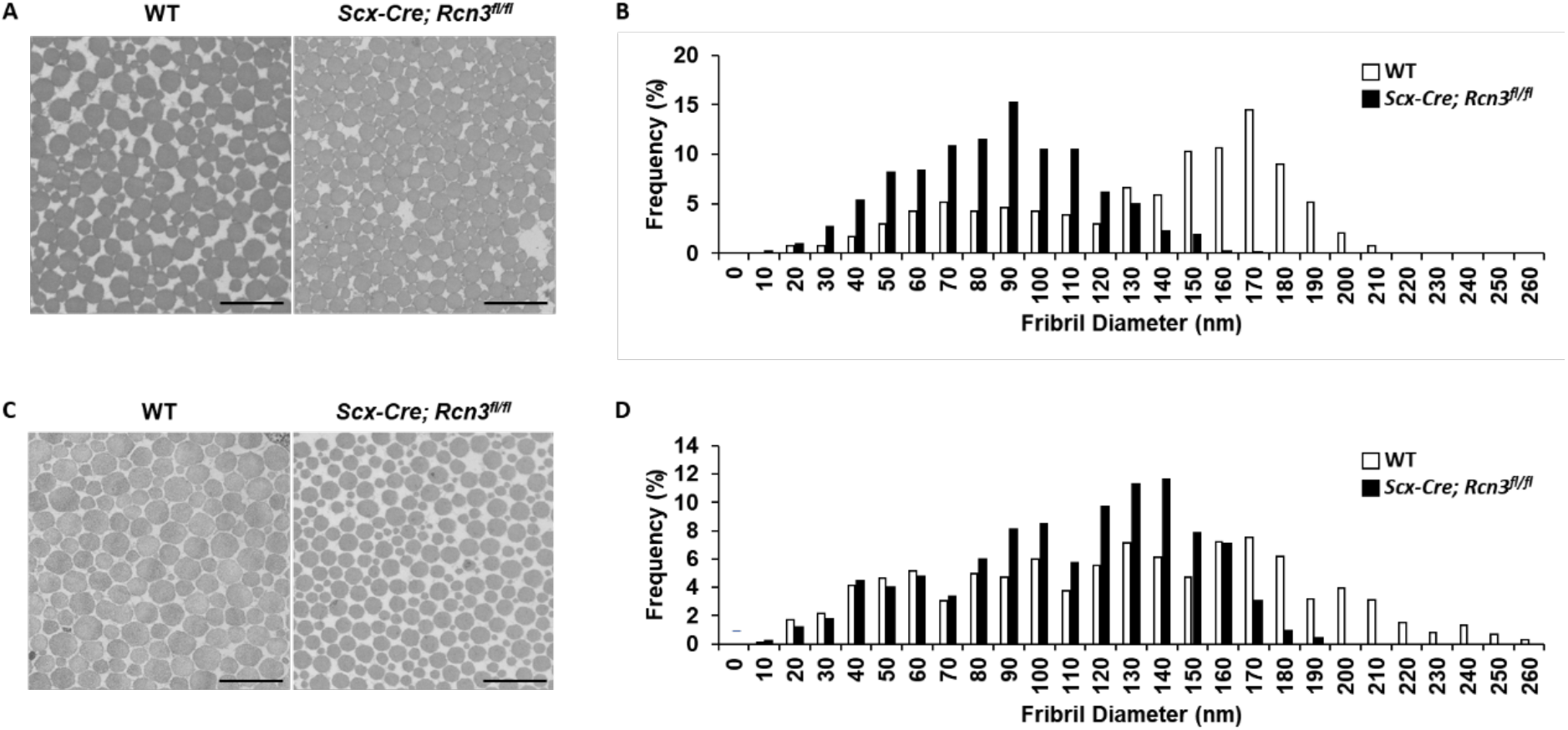
EM image of patellar (A) and Achilles (C) tendons and frequency graph of patellar (B) and Achilles (D) tendons of wild-type mice and *Scx-Cre; Rcn3*^*fl/fl*^ littermates at P60. (Scale bar indicates 500nm (A and C), n=3)

### Loss of Rcn3 increased collagen modification

Collagen post-translational modifications and cross-linking analysis were investigated in *Scx-Cre; Rcn3*^*fl/fl*^ mice at one month of age. The modification state of several key residues was examined using peptide mass spectrometry. Notable findings include the C-telopeptide cross-linking lysine from α1(I) chain, which was slightly over-hydroxylated in *Scx-Cre; Rcn3*^*fl/fl*^ patellar (20% more hydroxylated) and Achilles (9% more hydroxylated) tendons compared to wild-type mice (Table 1A). These data support that partial telopeptide lysine hydroxylation in tendon is being shifted to fuller hydroxylation in *Scx-Cre; Rcn3*^*fl/fl*^ mice. This is a useful mark of altered collagen cross-linking and is consistent with an increase in pyridinoline cross-links. Indeed, the cross-linking analysis revealed an increase in hydroxylysylpyridinoline in both tendons of the *Scx-Cre; Rcn3*^*fl/fl*^ mice. It should be noted that while hydroxylysylpyridinoline provides a useful measure of stable (irreversible) cross-links, it is not a measure of the total cross-links (Table 1B). The pattern of collagen over-modification was also observed across the sites of proline 3-hydroxylation in type I collagen in *Scx-Cre; Rcn3*^*fl/fl*^ Achilles and patellar tendons (Table 1). A function has yet to be discovered for this rare modification, but it has been suggested that 3-hydrxyproline can regulate the postnatal lateral fusion of smaller fibrils in developing tendons^23^.

**Table 1.** Collagen cross-link and mass spectral analysis to investigate the hydroxylation levels (A) of lysine and proline in type 1 collagen (n=2) and the collagen cross-linking levels (B) (* indicates *P*<0.05 between genotypes, n=1 for patellar and n=3 for Achilles tendon) from wild-type mice and *Scx-Cre; Rcn3*^*fl/fl*^ (tendon-specific Rcn3 loss-of-function model) littermates at P30.

| Hydroxylation analysis |  | Patellar tendon |  | Achilles tendon |  |
| --- | --- | --- | --- | --- | --- |
| Site |  | WT | *Scx-Cre; Rcn3<sup>fl/fl</sup>* | WT | *Scx-Cre; Rcn3<sup>fl/fl</sup>* |
| Hydroxylysine | $\alpha 1(I)$K87 | 100% | 100% | 100% | 100% |
| | $\alpha 2(I)$K87 | 100% | 100% | 100% | 100% |
| | $\alpha 1(I)$ C-telo Hyl | 45% | 65% | 39% | 48% |
| 3-Hydroxyproline | $\alpha 1(I)$P986 | 92% | 96% | 92% | 94% |
| | $\alpha 1(I)$P707 | 60% | 80% | 50% | 71% |
| | $\alpha 2(I)$P707 | 45% | 65% | 77% | 87% |

**Table 1.** Collagen cross-link and mass spectral analysis to investigate the hydroxylation levels (A) of lysine and proline in type 1 collagen (n=2) and the collagen cross-linking levels (B) (\* indicates $P<0.05$ between genotypes, n=1 for patellar and n=3 for Achilles tendon) from wild-type ...
| Cross-linking analysis |  | Patellar tendon |  | Achilles tendon |  |
| --- | --- | --- | --- | --- | --- |
|  |  | WT | *Scx-Cre; Rcn3<sup>fl/fl</sup>* | WT | *Scx-Cre; Rcn3<sup>fl/fl</sup>* |
| Hydroxylysylpyridinoline (mol/mol) |  | 0.25 | 0.36 | 0.09 | 0.17 * |

### Loss of Rcn3 caused abnormal tendon cell morphology

To investigate the cellular phenotype caused by the deletion of *Rcn3*, we analyzed tendon cell morphology using confocal microscopy after Zo1 and phalloidin double staining (Figure 4A). Tendon cell areas became largest at P30 and dramatically decreased at P60 in wild-type mice (Figure 4B, white bar). Interestingly, tendon cells in *Scx-Cre; Rcn3*^*fl/fl*^ mice exhibited increased cell area at P10 and P21 but significantly decreased cell areas at P30 compared to wild-type mice (Figure 4B). We also found changes in protrusion number in tendon cells of *Scx-Cre; Rcn3*^*fl/fl*^ mice. These protrusions from the cell body are critical for tendon cell interaction with adjacent cells and the ECM. The protrusion number number, in these transverse sections, decreased during postnatal tendon maturation in wild-type mice (Figure 4C, white bar). The *Scx-Cre; Rcn3*^*fl/fl*^ mice showed a significantly increased protrusion number compared with wild-type mice at all stages of postnatal tendon development (Figure 4C). Collectively, these results suggest that the function of Rcn3 is indispensable for the morphological maturation of tendon cells.

**Figure 4.**
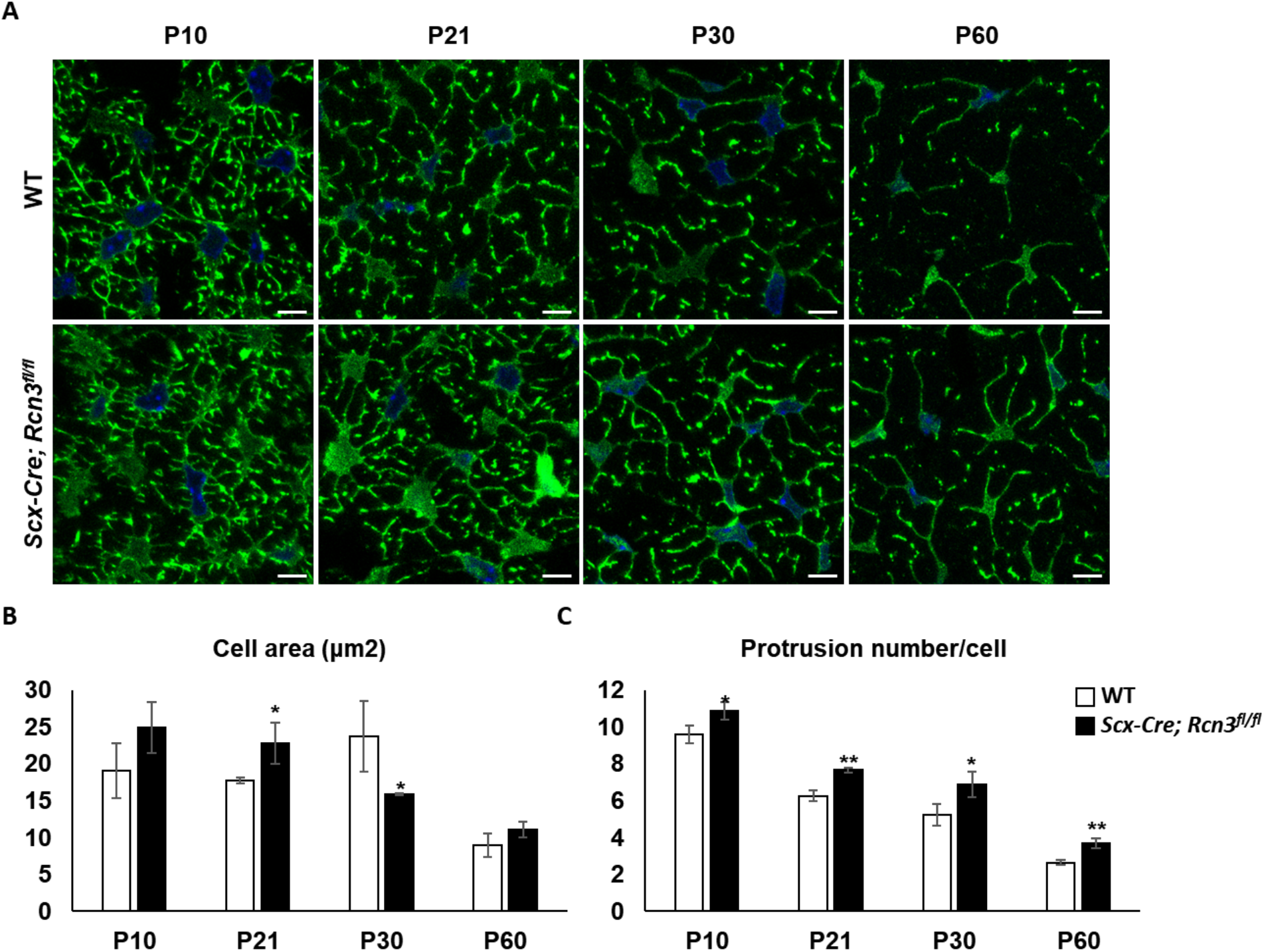
Immunofluorescence staining of ZO1 and phalloidin double staining on cross-sectional patellar tendon from wild-type mice and *Scx-Cre; Rcn3*^*fl/fl*^ (tendon-specific Rcn3 loss-of-function model) littermates during postnatal tendon maturation (A). Cell area (B) and protrusion number (C) of the patellar tendon. (Scale bar indicates 5µm (A), * indicates *P*<0.05, and ** indicates *P*<0.01 between genotypes, n=3).

### Loss of Rcn3 enhanced cellular maturation

To further understand the cellular phenotype at the molecular level, we performed qRT-PCR using Achilles tendon from wild-type mice and *Scx-Cre; Rcn3*^*fl/fl*^ mice. First, we confirmed the efficient deletion of Rcn3 in the Achilles tendon of *Scx-Cre; Rcn3*^*fl/fl*^ mice (Figure 5). The expression of tenogenic markers, such as *Scx, Mkx, Col1a1*, and *Tnmd*, were significantly increased in *Scx-Cre; Rcn3*^*fl/fl*^ mice compared with wild-type mice (Figure 5). These results suggest that the loss of Rcn3 enhanced the tendon cell maturation.

**Figure 5.**
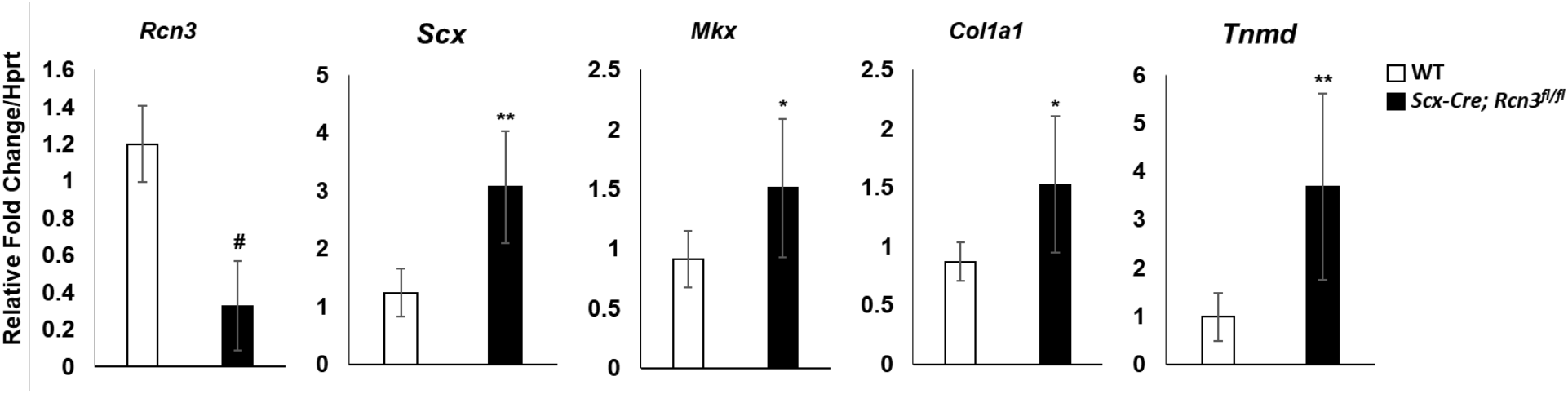
Quantitative real-time PCR of Achilles tendons from wild-type mice and *Scx-Cre; Rcn3*^*fl/fl*^ (tendon-specific Rcn3 loss-of-function model) littermates at P28 (C). (* indicates *P*<0.05, ** indicates *P*<0.01 and ^#^ indicates *P*<0.001 between genotypes, wild-type mice n=6, *Scx-Cre; Rcn3*^*fl/fl*^ mice n=5)

### Loss of Rcn3 decreased mechanical properties

To measure the functional outcome of Rcn3 loss in tendons, we performed uniaxial biomechanical testing of the Achilles tendon at two months of age. Consistent with our histological analysis, cross-sectional area (CSA) was significantly reduced in *Scx-Cre; Rcn3*^*fl/fl*^ tendons (Figure 6A). Concomitantly, stiffness and failure load were also significantly lower, suggesting a loss of structural capability in the *Scx-Cre; Rcn3*^*fl/fl*^ mice (Figure 6B and 6C). No difference was observed in percent stress relaxation between the two groups, suggesting that Rcn3 does not perturb the viscoelastic response of the Achilles tendon (Figure 6D). In addition, no differences were observed in failure stress or tissue modulus, which indicates that loss of Rcn3 did not cause an inherent material change in the tendon tissue (Figure 6E and 6F). These data demonstrate that reduced mechanical properties caused by loss of Rcn3 are due to decreased structural properties but not material properties.

**Figure 6.**
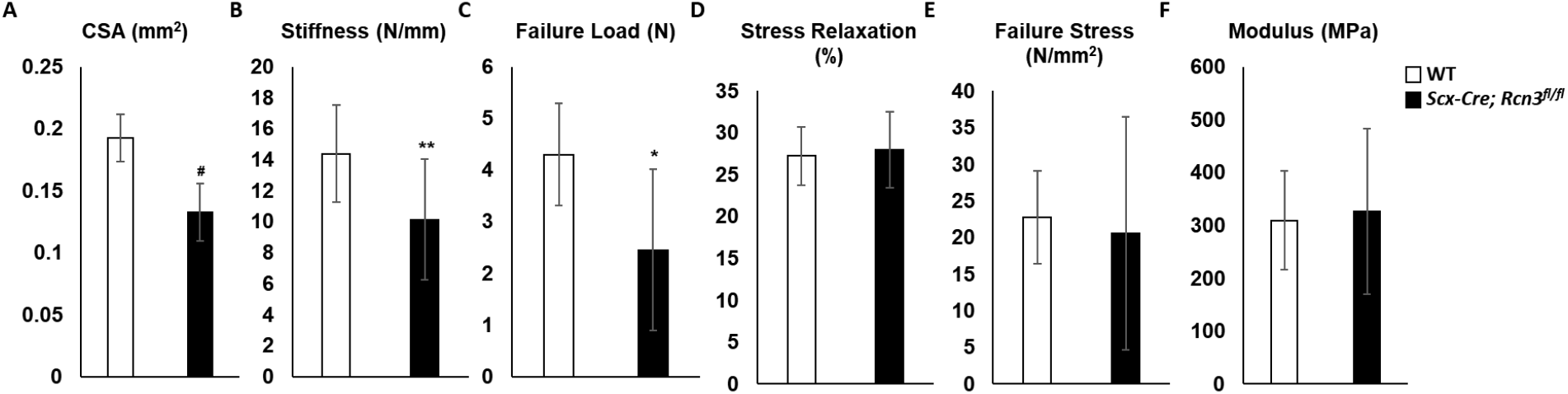
Mechanical property, cross-sectional area (A), stiffness (B), failure load (C), stress relaxation (D), failure stress (E), and modulus (F) of Achilles tendons from wild-type mice and *Scx-Cre; Rcn3*^*fl/fl*^ (tendon-specific Rcn3 loss-of-function model) littermates at P60. (* indicates *P*<0.05, ** indicates *P*<0.01, and ^#^ indicates *P*<0.001 between genotypes, n=7)

## Discussion

We investigated the function of Rcn3 in postnatal tendon development by characterizing a tendon-specific Rcn3 knockout mouse (*Scx-Cre; Rcn3*^*fl/fl*^) model. The *Scx-Cre; Rcn3*^*fl/fl*^ mice exhibited impaired postnatal tendon development, including decreased tendon thickness, increased cellularity, impaired tenocyte maturation, and reduced collagen fibril diameter with altered collagen modification. Our study identified the novel function of Rcn3 in tendon and provides genetic approaches that can be used to investigate the underlying mechanisms regulating cellular and matrix maturation in tendon.

Rcn3 is known as an ER protein regulating the secretion of proteins, but its critical molecular function is not clear. Our study showed that Rcn3 deficiency caused impaired collagen fibrillogenesis with over-modification of collagen. These results suggest that Rcn3 is a critical regulator of collagen fibrillogenesis via regulation of collagen modification. However, the precise molecular mechanism by which Rcn3 regulates collagen modification is still not clear. One possibility is that Rcn3, as a chaperon protein, stabilizes collagen for proper modification. Previous studies also suggested that Rcn3 functions as a chaperon protein^27,28^. The other possibility is that Rcn3 mediates the binding of collagen modification enzymes to collagen molecules. Further molecular analyses will be required to elucidate the precise function of Rcn3 in collagen modification in tendon.

To measure morphological changes of tendon cells at a single-cell level, we developed a novel confocal imaging method (Figure 4). We found that the protrusion number was decreased throughout the postnatal tendon development in wild-type mice. Interestingly, this result is different from the current finding using serial block face-scanning electron microscopy (SBF-SEM)^12^. Kalson et al. showed that a similar number of protrusions contacting adjacent cells between the tendons of newborn and six-week-old mice. There are several variations between our and Kalson’s study. First, we have analyzed the patellar tendon, but they analyzed the tail tendon. Second, the age is different. We have analyzed age from P10 to P60. They analyzed newborn and six-week-old mice. These age and tendon types could cause different results of tendon phenotype in each study. Further studies analyzing various tendon types and ages using our confocal imaging technique and SBF-SEM will be necessary to fully understand the morphological maturation of tenocytes.

During normal tendon development, the average cross-sectional cell area became largest at P30 in wildtype (Figure 4). However, *Scx-Cre; Rcn3*^*fl/fl*^ mice showed a prematurely increased cell area with increased cell area at P10 and P21, while significantly decreased cell area at P30 compared to wildtype. These results suggest that loss of Rcn3 causes abnormal organization of tenocytes. These cell shape changes correspond with the molecular analysis (Figure 5) that shows increased expression of tenogenic markers in Rcn3 deficient mice. Based on these data, we concluded that Rcn3 negatively regulates the maturation of tenocytes. Investigating the mechanism by which Rcn3 regulates tenocyte maturation will be an interesting future direction of this study.

*Scx-Cre; Rcn3*^*fl/fl*^ mice displayed an increase in post-translational modifications and cross-linking in type I collagen. Specifically, *Scx-Cre; Rcn3*^*fl/fl*^ mice showed an increase in telopeptide lysine hydroxylation as well as hydroxylysylpyridinoline in both patellar and Achilles tendons. Hydroxylysylpyridinoline is a stable collagen cross-link, typically found in adult bone and tendon, that is known to increase in concentration as these tissues mature. Besides, *Scx-Cre; Rcn3*^*fl/fl*^ mice exhibited over-hydroxylated proline 3-hydroxylation in type I collagen. The function of this rare modification is not clear, but it has been suggested that 3-hydrxyproline can regulate the postnatal lateral fusion of smaller fibrils in developing tendons^23^. Based on these collagen modification data, we carefully suggest that the reduced collagen fibril diameter in *Scx-Cre; Rcn3*^*fl/fl*^ mice is due to the inhibition of lateral fusion caused by over-modification on type I collagen.

We observed the higher deletion efficiency of Rcn3 in the patellar tendon than the Achilles tendon (Figure 1B). Consistent with this result, the overall phenotypes of *Scx-Cre; Rcn3*^*fl/fl*^ mice were more pronounced in patellar tendons. These data support the dosage-dependent function of Rcn3 in postnatal tendon development. The different deletion efficiency could be due to the different expression levels of CRE recombinase in *Scx-Cre* line and/or the accessibility of the endogenous *Rcn3* locus to recombination in these different tendons. Further studies will be required to determine the expression level or the efficiency of CRE recombinase in the different tendons of *Scx-Cre* mouse line.

In conclusion, our results uncover the novel function of Rcn3 in postnatal tendon development. This study will be the basis of future mechanistic studies for tenocyte maturation, collagen modification, and fibrillogenesis by providing useful mouse genetic models.

## Methods

### Animals

Animal care and experiments were performed in accordance with the guidelines issued by the Institutional Animal Care and Use Committee of the University of Pennsylvania (Philadelphia, Pennsylvania, USA). The *Scx-Cre* mouse line was previously described^33^. Mouse sperms carrying Knockout-first *Rcn3* allele (*Rcn3* ^*tm1a(EUCOMM)Hmgu*^) were obtained from the Knockout Mouse Programme (KOMO), and mouse carrying *Rcn3* ^*tm1a(EUCOMM)Hmgu*^ was generated by in vitro fertilization. *Rcn3* ^*tm1a(EUCOMM)Hmgu*^ mice were crossed with *Rosa26*-*Flippase* (*Flp*) mice to delete β-gal and the neo cassette in order to generate the *Rcn3* conditional knockout allele (*Rcn3*^*fl/fl*^).

### Immunohistochemistry

For immunohistochemisty, mouse hindlimbs were collected during postnatal tendon maturation from P10 to P60 and fixed in 10% neutral buffered formalin overnight at 4°C. Samples were paraffin-embedded and sectioned at 6 μm, following decalcification in 10% (w/v) EDTA (pH 7.4) for 2 weeks with daily solution changes. The paraffin sections of the patellar and Achilles tendon incubated with antibodies against Rcn3 (Abcam, ab204178) at 4°C overnight. The sections were then incubated in anti-rabbit-biotin antibody (Jackson Immuno, 711-065-152) for 30 min at RT, rinsed with PBS, and incubated with streptavidin-HRP (Jackson Immuno, 016-030-084) for 30 min at RT. Bounded antibodies were detected using NovaRed substrate (Vector Laboratories, SK-4800).

### Histological analyses

Mouse hindlimbs were collected from specific stages and fixed in 10% neutral buffered formalin overnight at 4°C. Samples were paraffin-embedded and sectioned at 6 μm, following decalcification in 10% (w/v) EDTA (pH 7.4) for 2 weeks with daily solution changes. The paraffin sections of the patellar and Achilles tendons were used for Hematoxylin and Eosin (H&E). Histological evaluation was blindly performed by 2 independent individuals.

### Collagen EM analysis

Mouse hindlimbs were fixed in freshly prepared 1.5% glutaraldehyde/1.5% paraformaldehyde (Tousimis) with 0.05% tannic acid (Sigma) in DPBS at 4 °C with agitation overnight. The following day, dissect the relevant tendons out of the limb in PBS as follows: for the Achilles, take the mid-tendons that lies below the myotendinous junction and above the enthesis (∼1-1.5 mm in length); and for the patellar tendon, take the complete mid-tendon between the patella and tibial enthesis. Following dissection, samples were post-fixed in 1% OsO4, rinsed in DMEM and dehydrated in a graded series of ethanol to 100%. Samples were then rinsed in propylene oxide, infiltrated in Spurrs epoxy, and polymerized at 70 °C overnight. TEM images were acquired using a FEI G20 TEM at multiple magnifications to visualize transverse sections of collagen fibrils. Collagen fibril diameter was measured using ImageJ.

### Collagen extraction

Intact type I collagen was solubilized from mouse patellar and Achilles tendons by acid extraction in 3% acetic acid for 24h at 4°C. Acid-extracted collagen α-chains were resolved by SDS-PAGE and stained with Coomassie Blue R-250.

### Collagen cross-linking analysis

Collagen pyridinoline cross-link content was determined by fluorescence monitoring with reverse-phase HPLC. Pyridinoline cross-links were analyzed in mouse tendons by HPLC after acid hydrolysis in 6M HCl for 24h at 108°C. Dried samples were dissolved in 1% (v/v) n-heptafluorobutyric acid for quantitation of hydroxylysyl pyridinoline (HP) by reverse-phase HPLC and fluorescence monitoring as previously described^34^.

### Mass spectrometry

Collagen α-chains were cut from SDS-PAGE gels and subjected to in-gel trypsin digestion as previously described^24^. Electrospray mass spectrometry was carried out on the trypsin-digested peptides using an LTQ XL linear quadrupole ion-trap mass spectrometer equipped with in-line Accela 1250 liquid chromatography and automated sample injection (ThermoFisher Scientific). Proteome Discoverer software (ThermoFisher Scientific) was used for peptide identification. Tryptic peptides were also identified manually by calculating the possible MS/MS ions and matching these to the actual MS/MS spectrum using Thermo Xcalibur software. Differences in post-translational modifications were determined manually by averaging the full scan MS over several LCMS minutes to include all the post-translational variations of a given peptide. Protein sequences used for MS analysis were obtained from the Ensembl genome database.

### Immunofluorescent staining

For immunofluorescent staining, mouse hindlimbs were collected during postnatal tendon maturation from P10 to P60 and fixed in 4% PFA overnight at 4°C. Samples were OCT compound embedded and sectioned at 10 μm, following decalcification in 10% (w/v) EDTA (pH 7.4) for 10 days with daily solution changes. The frozen sections of the patellar tendon blocked with blocking solution (Thermo, 8120) for 1hr RT, followed by incubation at 4°C overnight with antibodies against ZO1 (Fisher, 61-7300) and then incubated for 1hr at RT with Phalloidin Alexa Fluor™ 488 conjugate. Finally, sections were mounted in antifade solution with DAPI (Invitrogen, P36935).

### Primary tendon cell culture

For primary tendon cell culture, the tail was collected form P25 in PBS on ice. The skin was removed and then pull out the tendon from the tail. Tail tendons were incubated for 1 hr at 37°C in 2mg/ml collagenase (Sigma, C0130-1G). Tail tendon and cells which were collected by centrifugation were plated on culture dishes. Cells were maintained in growth media (αMEM, 20% FBS, 2mM L-glutamine).

### qRT-PCR

RNA was extracted with Trizol and Direct-zol kit (Zymo Research, R2060) from primary tendon cells. And cDNA was synthesized from 1μg RNA using iScript Reverse Transcription (Bio-rad, 1708841). qRT-PCR was performed using Fast SYBR Green Master Master Mix (Pec, 4385612). The sequences of primers are listed in Table 2.

**Table 2.**
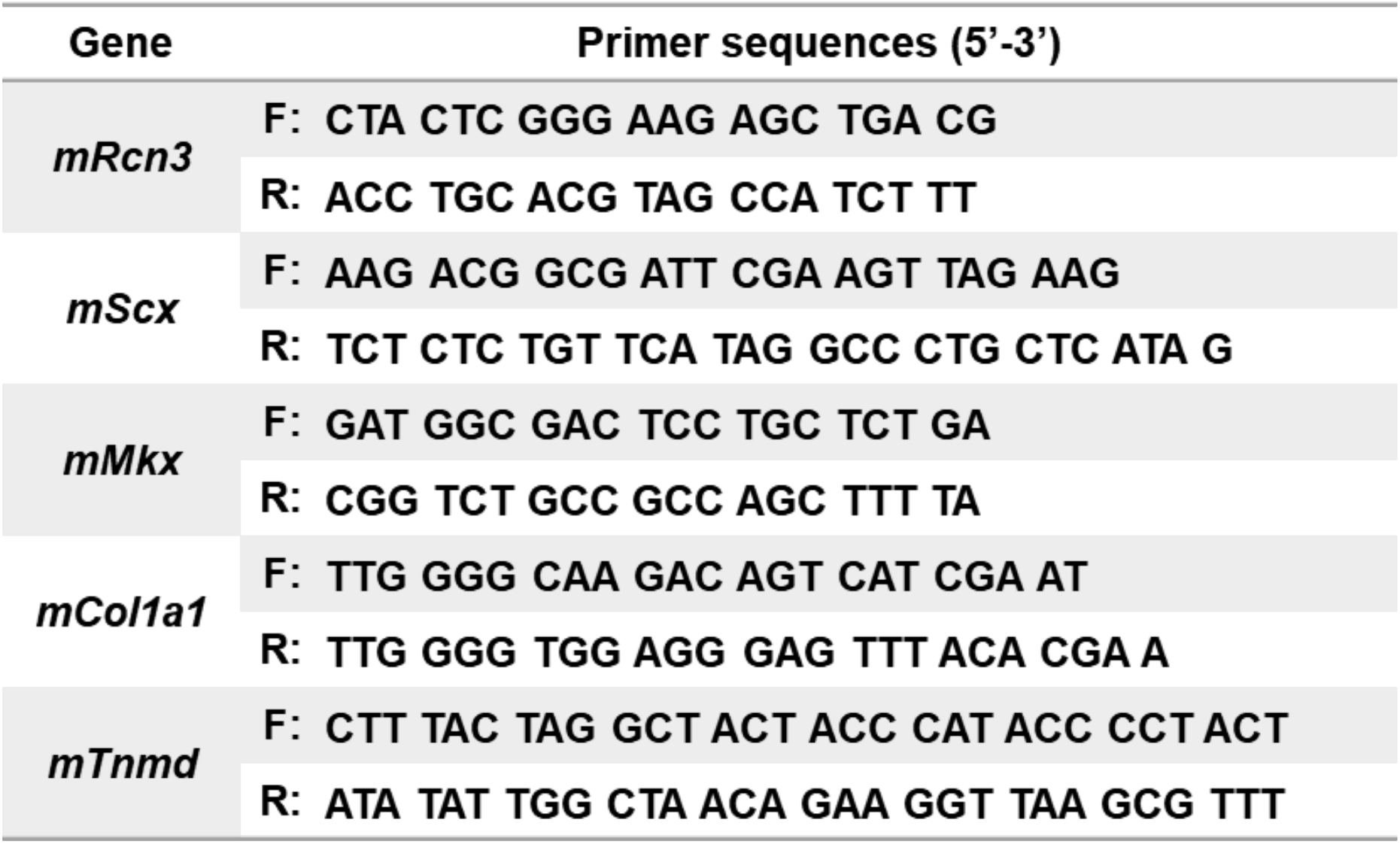
Primer sequences for the quantitative real-time PCR.

### Uniaxial biomechanical test

All mice assigned for mechanical testing were frozen at -20C untill the day of testing. Mice were thawed at room temperature, and calcaneal bone – Achilles tendon – muscle complexes were grossly dissected. Under microscope guidance, all extraneous soft tissues and muscles were finely dissected with care to ensure no damage to the tendinous tissue. Following dissection, a custom laser device was used to measure the cross-sectional area (CSA) of the Achilles tendon. The myotendinous junction was sandwiched between two sandpaper tabs with cyanoacrylate glue to prevent slippage. The calcaneal bone was gripped with a custom fixture, and the construct was mounted onto a material testing machine (Instron 5542, Instron Inc., Norwood, MA). All testing was conducted in phosphate buffered saline bath at room temperature. Each sample was preloaded to 0.02N followed by 10 cycles of preconditioning between 0.02 to 0.04N. After a resting period of 300 seconds at 0N, the sample underwent stress relaxation after a ramp to 5% strain (assuming a gauge length of 5mm) for 600 seconds. This was followed by a rest of 60 seconds at zero load. Finally, the sample was quasi-statically ramped to failure at a strain rate of 0.03%/s. All data were collected at 100Hz. Ensuing force-displacement curves were analyzed to obtain failure load (N) and tissue stiffness (N/mm, defined as the slope of the linear region). Cross-sectional areas (mm2) and gauge length (mm) values were used to obtain stress-strain curves for each sample. Modulus (N/mm2) was calculated as the slope of the linear region of the stress-strain curve and failure stress (N/mm2) as the maximum stress value observed. Stress relaxation (%) was defined as the percent change in stress between the peak and equilibrium stress during the stress relaxation period of 600 seconds.

### Statistical analysis

Results are expressed as mean ± SD. At least three mice per group were analyzed. Differences between values were analyzed by Student’s t-test. P<0.05 is considered significant.

## Author contributions

Study conception and design: KSJ, NAD, NRP; Acquisition of data: NRP, SS, DRK, ST, DMH, MA; Analysis and interpretation of data: NRP, DMH, DRK, LJS, NAD, KSJ; Drafting of the manuscript: KSJ, NRP, DMH, DRK

## Acknowledgments

We would like to thank Penn Center for Musculoskeletal Disorders (PCMD) Histology and Biomechanics Core for technical assistance with the histology and biomechanical testing, respectively. We also thank Cell and Developmental Biology (CDB) Microscopy Core for technical assistance with confocal microscopy. We thank Dr. Ronen Schweitzer for providing *Scx-Cre* mouse line.

